# Comparing freshwater eDNA with conventional survey methods to detect vertebrate biodiversity in a Malagasy tropical rainforest

**DOI:** 10.1101/2025.08.28.672370

**Authors:** Christina Biggs, Cosmo Le Breton

## Abstract

Madagascar is globally recognized for its exceptional biodiversity and endemism yet remains severely under-sampled for many taxa. We evaluate the effectiveness of environmental DNA (eDNA) metabarcoding as a rapid, non-invasive method for surveying vertebrate biodiversity in the Makira Forest Protected Area (MFPA), comparing its performance to conventional surveys across a range of vertebrate taxa. eDNA sampling detected 158 OTUs across 17 sites, outperforming conventional surveys in birds (*p* = 0.000107), mammals (*p =* 0.000718) amphibians (*p* = 0.000725), reptiles (*p =* 0.000877) and ray-finned fish (*p* = 0.00788). We further examined the result for ray-finned fish by modelling species-accumulation curve asymptotes which supported this result. However, significant primer bias and amplification inconsistencies were observed, particularly in amphibian detections, and a high proportion of OTUs could not be resolved to species level due to a combination of taxonomic gaps in reference databases, DNA degradation affecting sequence quality or length, and marker-specific limitations. We demonstrate, using species accumulation curves of our eDNA data and inventoried taxa lists of the MFPA, that the widespread practice of considering only resolved OTUs, rather than all unresolved OTUs, significantly underestimates biodiversity. This study underscores both the promise and current limitations of eDNA as a tool for tropical biodiversity assessment, specifically when compared to conventional sampling techniques. Combining eDNA and conventional survey methods increased species detection rates by 66.4% in comparison to the use of visual surveys alone, highlighting their complementary strengths and reinforcing the value of integrated monitoring strategies for biodiversity-rich, data-deficient regions.

## INTRODUCTION

Madagascar is renowned for both its high species richness and high rates of endemism, shaped by the island’s complex geological history and continental isolation [1]. Classified as one of the world’s 17 megadiverse countries, Madagascar is one of the world’s eight ‘hottest’ biodiversity hotspots [2], with over 300 species of birds [3], 150 species of mammals [4], 600 species of herpetofauna [5], and 100 species of freshwater fish [6]. Madagascar is also unique due to its extremely high levels of endemism, with 50% of bird species, and 98% of reptiles, amphibians and mammals being endemic to the island [7]. While Madagascar has long been recognized as a conservation priority [8], acute deforestation has destroyed over 40% of its original forest cover since the 1950s, resulting in severe biodiversity decline and ecosystem instability [9].

Makira Forest Protected Area (MFPA), located on the north-east coast of Madagascar, covers more than 372,000 ha of one of the largest remaining expanses of intact tropical rainforest on the island. Formally established in 2012, the MFPA forms part of the 98 strategic protected areas in Madagascar, providing connectivity between the Masoala and Marojejy National Parks. The MFPA is estimated to contain 316 species: 112 bird species, 58 species of mammals including 17 lemurs [10], 68 amphibians [11], 60 reptiles [12], and 18 species of freshwater fish [13]. The MFPA also remains a top conservation priority due to the number of threatened species it contains, 46% of the total Malagasy Red Listed species according to IUCN [14]. Monitoring biodiversity in these forests is essential for assessing conservation priorities and guiding protection efforts, yet conventional field surveys remain logistically challenging, time-intensive, and expensive, with limited capacity to detect rare or cryptic species.

As a form of non-invasive, high throughput ecological surveillance, eDNA metabarcoding analysis has revolutionized the field of biodiversity research by allowing the identification of invasive, cryptic (both visually and genetically) and undescribed species, with broad applications to understanding ecosystems[15-19]. The detection of DNA persisting in ecosystems can be used to provide information on species, populations, and communities, representing a significant breakthrough as a tool for ecologists and conservation policy making [20]. The benefits over other non-invasive sampling methods; such as cost effectiveness, time efficiency, scalability, and both comprehensiveness and equitability could be significant [21]. However, while the technical capabilities of eDNA are now well-established, less attention has been paid to how these tools can be best deployed to generate accurate ecological baselines datasets essential to informing conservation priorities, allocating resources, and evaluating policy interventions [22, 23]. In regions where biodiversity is both exceptional and understudied, establishing such baselines is critical for preventing extinctions and guiding protected area management.

Despite Madagascar representing a clear conservation priority, it remains a globally significant darkspot of biodiversity across multiple taxa [24, 25]; a point of serious concern for conservation policy in the face of anthropogenic pressures. Although previous research efforts in the MFPA focused on specific taxonomic groups such as carnivores and palms, significant data deficiencies of the park’s overall biodiversity persist. In this eDNA metabarcoding study, we compare the efficacy of tetrapod eDNA analysis with conventional surveys completed by specialist taxonomists within the MFPA to produce updated biodiversity baselines[26]. This study was completed as part of the broader “Search for Lost and Undescribed Species: A rapid, multi-taxon biodiversity inventory of Makira Natural Park, Madagascar,” conducted from August 29th to September 9th, 2023. This study aims to evaluate and compare the effectiveness of eDNA metabarcoding and conventional surveys for assessing multi-taxa vertebrate biodiversity in a tropical rainforest, while identifying methodological limitations and opportunities to improve future biodiversity inventories.

## METHODS

All fieldwork activities were conducted in accordance with the research permits issued by the Malagasy Ministry of Environment and Sustainable Development, outlined in the research permit 273/23/MEDD/SG/DGGE/DAPRNE/SCBE.Re of 09/08/2023, with specimen exports authorized by permit number 39 IN-EA10/MG23 of 10/13/2023.

### 2.1 Survey Site

All surveys and sampling were conducted from August 29 to September 9, 2023, in the Sahamatreha forest within MFPA, Madagascar (S 15°04’15.2”, E 49°34’48.3”), 44 km northwest of Maroantsetra, at altitudes between 250–700 m. Designated as ‘Af’ under the Köppen climate classification system, the area is characterized by tropical rainforest with consistently high temperatures and significant rainfall throughout the year, with no distinct dry season and extensive humid forest [27]. Locations and survey distances of all survey techniques were recorded using Garmin inReach Explorer+.

### 2.2 eDNA Analysis

#### 2.2.3 Sampling

Field sampling followed the NatureMetrics Standard Aquatic eDNA Kit protocol [28].The 20 eDNA sampling locations were selected in river, stream and lake environments (precise locations being determined by water flow dynamics) [29] (Figure 1). They targeted aquatic habitats along the drainage network within the broader Sahamatreha survey area. Conventional surveys (birds, mammals, fish, herpetofauna) were conducted within the same landscape but were not spatially paired to individual eDNA sampling locations. At each site, three independent samples were collected from the left bank, centre, and right bank. Sampling progressed upstream, starting from the most downstream location, with all waterways accessed on foot. Gloves were worn, and sediment disturbance was minimized. Water was collected at the surface using a 5 L dedicated sampling bag, with the sampler downstream to avoid contamination. The bag was sealed and shaken for 30 seconds. For sediment-heavy samples, water was allowed to settle for one minute. Each sample was subsequently filtered in repeated 60 mL syringe draws (6–50 draws per sample), corresponding to total filtered volumes ranging from 380 mL to 3,000 mL, until the filter clogged or the entire sample was processed. Air was then used to expel residual water. A preservative syringe was attached to the filter inlet, and Longmire’s solution was slowly injected. Both ends of the filter were capped with Luer Lock caps, and the filter was sealed in a specimen bag. Each sample’s Barcode, ID, date, GPS coordinates, volume filtered, habitat characteristics, and sampler information were logged. A field negative control was included at the site. A field negative control was included at each site and consisted of 900 mL of commercially purified bottled water, which was processed identically to field samples, including filtration, preservation, and handling steps. Site photographs and observations on flow, sedimentation, and weather were also recorded.

**Figure 1:**
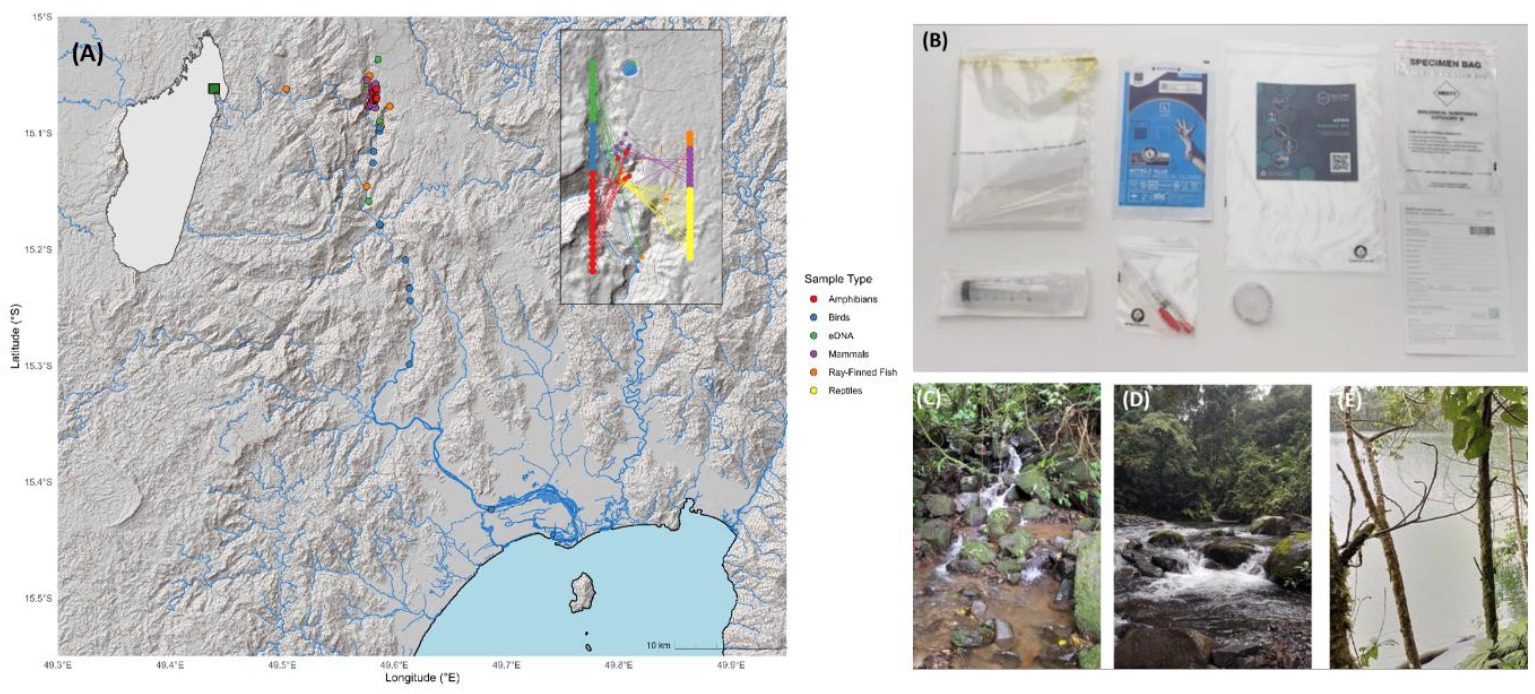
Survey design and preparation of eDNA sampling in the MFPA. Map of all survey sites (A) with clustered central sites shown in inset (R) and map of Madagascar inset (L). eDNA sampling equipment (B). Sample sites (C-E) showing range of lotic ecosystems sampled. MAK#2 (C), MAK#8 (D), MAK#19 (E).

#### 2.2.2 OTU Extraction & Amplification

Two sites were removed from subsequent analysis due to failed DNA extraction and sequencing (MAK17, MAK20). Samples were processed in dedicated clean rooms for eDNA work at NatureMetrics in Class II biosafety cabinets, with workstations decontaminated using chemical disinfectant and UV irradiation before and after use. Samples were collected on 0.8 µm polyethersulfone (PES) filters, with modified Longmire’s solution added to the filter housing for preservation prior to extraction. DNA was extracted from enclosed 0.8 µm PES filters using a DNeasy Blood and Tissue Kit (Qiagen, 1001 Marshall St, Redwood City, CA 94063; https://www.qiagen.com/us) following the Spens et al. enclosed filter protocol [30], with proteinase K added directly to the filter housing to minimize contamination risk. Extracts were subsequently purified using the DNeasy PowerClean Pro Cleanup Kit (Qiagen, 1001 Marshall St, Redwood City, CA 94063; https://www.qiagen.com/us) to remove PCR inhibitors. Negative controls consisting of molecular-grade water were processed with each batch through extraction and inhibitor-removal steps. DNA concentration was measured using a Qubit fluorometer with the Qubit dsDNA Broad Range assay kit (Thermo Fisher Scientific, 340 El Pueblo Rd, Scotts Valley, CA 95066; https://www.thermofisher.com/us). Replicate PCRs targeted the mitochondrial 12S rRNA gene of freshwater vertebrates using primers F1 (5’-ACTGGGATTAGATACCCC-3’) and R1 (5’-TAGAACAGGCTCCTCTAG-3’), generating an amplicon of approximately 106 bp, and the mitochondrial 16S rRNA gene using mammal-specific primers 16Smam1 (5’-CGGTTGGGGTGACCTCGGA-3’) and 16Smam2 (5’-GCTGTTATCCCTAGGGTAACT-3’), generating an amplicon of approximately 95 bp [31]. A blocking primer minimized non-target human DNA amplification (12S_V5_blkhum; [32]). Positive and negative controls (synthetic sequences and PCR-grade water) accompanied each plate. Amplification was visualized using gel electrophoresis.

#### 2.2.3 Sequencing

Following first-round PCR of 12S and 16S amplicons, products were purified using magnetic beads and indexed in a second PCR using Illumina dual-index primers according to the Illumina 16S Metagenomic Sequencing Library Preparation protocol (Illumina, Inc., 5200 Illumina Way, San Diego, CA 92122; https://www.illumina.com). Indexed libraries were then pooled and sequenced on an Illumina MiSeq platform using a V3 600-cycle kit (2 × 300 bp paired-end sequencing).Demultiplexing used bcl2fastq based on index tags. Paired-end reads were merged with USEARCH [33], requiring 80% overlap agreement. Primers were trimmed with cutadapt [34], and sequences were quality-filtered with USEARCH (error rate ≤0.01). Unique sequences were dereplicated and denoised with UNOISE to identify zOTUs [35]. Taxonomic assignments were made via blastn searches against the NCBI nucleotide database, with thresholds of 99%, 97%, and 95% for species, genus, and higher-level identifications [36, 37]. Assignments were cross-checked with GBIF occurrence data to ensure local relevance, adjusting taxonomy as necessary [38]. zOTUs were clustered into OTUs using USEARCH UPARSE and custom pipelines to reduce intra-specific variants while preventing over-clustering [39]. Chimeric sequences were removed during the UNOISE denoising step using standard USEARCH procedures [32], and OTU tables were generated at a 97% identity threshold. OTUs were annotated with IUCN Red List status and GRIIS invasive status using the rredlist interface [40]. Low-abundance OTUs (<20 reads per sample) and those assigned to humans or domesticated mammals were excluded. The minimum read threshold was applied as part of the standard NatureMetrics bioinformatics pipeline to reduce the likelihood of spurious detections arising from sequencing or PCR artefacts.

### 2.3 Ornithological Surveys

A range of standard ornithological survey techniques were used. Data collection coincided with the pre-breeding period for most Malagasy forest birds and included direct observations using eBird Mobile, Cornell Lab of Ornitholog [41], and mist-netting (Figure 2). Observers completed point counts along non-standardized transects from 6:00–11:00 when birds were most vocally active and least mobile, for seven days. This reduced the chance of repeated contact [42, 43].Twenty mist nets (5 of 12m and 15 of 6m) were installed near streams within 10–30 m of the watercourse and dense vegetation [44]. Nets were deployed from 5:30–17:30 daily, monitored every 30 minutes, and captured birds were photographed, weighed, banded with numbered aluminium rings, and released.

**Figure 2:**
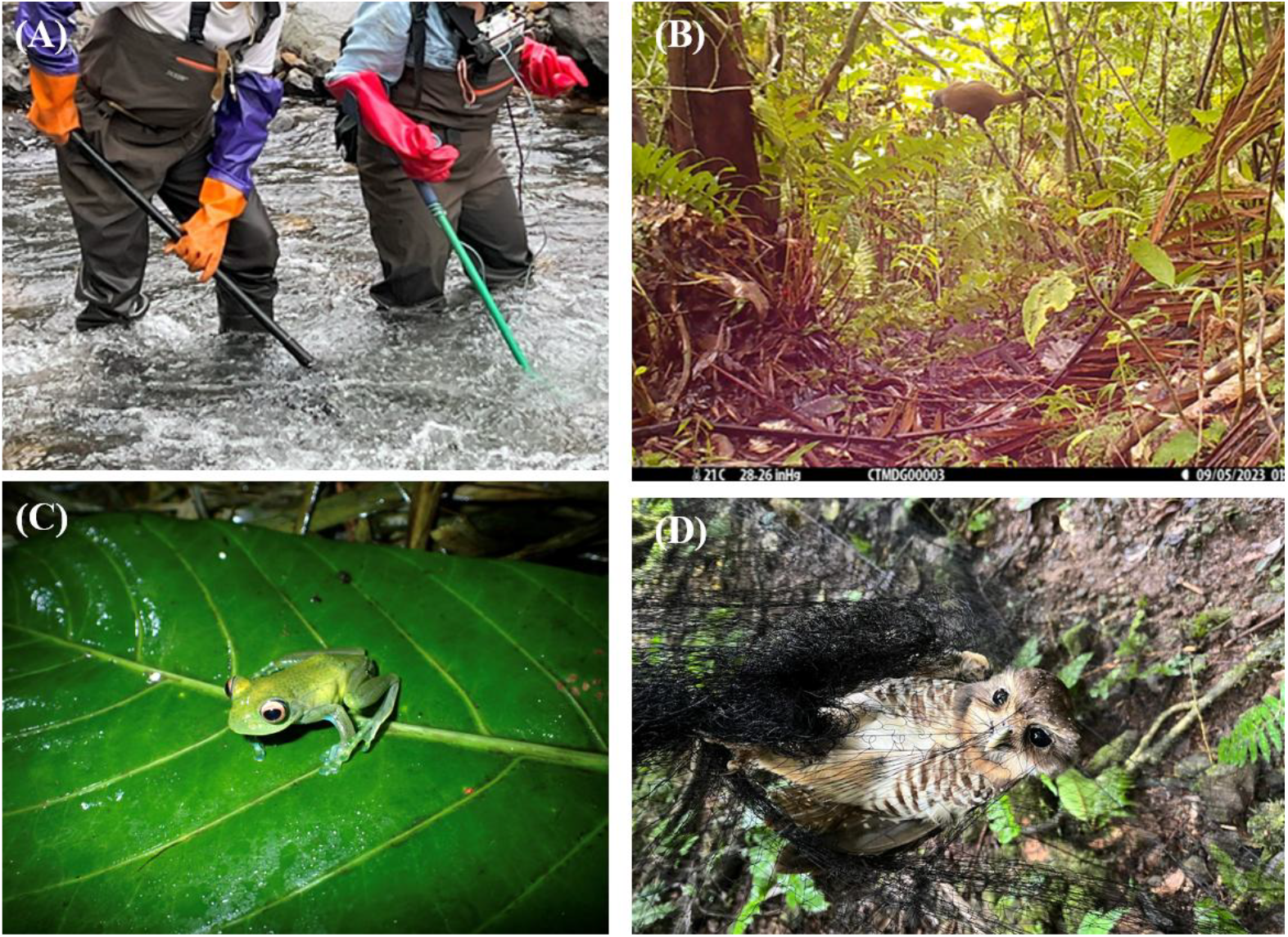
Selection of conventional survey techniques employed. Electrofishing (A), camera trapping of *Eulemur albifrons* (f.) (B), herpetofaunal transect identification of *Boophis englaenderi* (C); *Athene superciliaris* in a mist net (D).

Bird calls and photographs were recorded for threatened or notable species and uploaded to the eBird and Macaulay Library platforms.

### 2.4 Mammals Surveys

Mammal surveys were conducted using a combination of diurnal and nocturnal Visual Encounter Surveys (VES) along line transects as well as systematic searches, following methods adapted from Goodman et al. (1997) [45] and tailored for the terrain and target taxa. The primary transect was a pre-existing footpath that followed the altitudinal gradient of a canyon. Transects were surveyed twice daily 07:30–11:00 and 14:30–17:00 and again at night between 18:00–21:00. Nocturnal surveys utilized low-intensity torches to detect eyeshine, particularly for lemurs. Observers recorded all visual and auditory detections, including species ID, number of individuals, behavior, distance, height, and GPS location. In addition to transects, systematic searches were carried out in habitat features likely to harbor lemurs, such as areas with abundant fruiting trees, signs of feeding, or recent vocalizations using flexible routes based on topographic and ecological cues. Mist netting was conducted at two sites for *Chiroptera spp*. detection. Nets were set from 16:00–21:00 and 3:00–5:30.

Camera trapping was employed as a complementary method to document cryptic or disturbance-sensitive terrestrial and arboreal mammals that are difficult to detect using visual encounter surveys. A total of ten camera traps were operated continuously for five consecutive days (2–7 September 2023) along an altitudinal gradient between 200 and 760 m above sea level, primarily along existing trails and ridgelines near the main camp. Browning Dark Ops cameras were configured to record motion-triggered video sequences, with one-minute recordings during daylight hours and 20-second recordings at night. Camera trap placement was non-invasive and did not involve baiting.

### 2.5 Ichthyological Surveys

Ichthyological survey sampling sites were chosen based on accessibility, proximity to the Sahamatreha River and its tributaries, and historical records of target species. Additional considerations included habitat diversity, hydrological features, and input from local fishers on species presence. Seine netting and repurposed shrimp traps were deployed in areas of low depth and velocity. A microprocessor SUM unit (Electro Fisher Company Co. Ltd., 15 Bialostoczek St, Bialystock, BIA 15869, Poland; https://www.electro-fisher.net) was used in fluvial stretches of high turbidity, high velocity and areas of deeper bathymetry to induce “electronarcosis” and enabling capture of immobilized fish (Figure 2) [46].

### 2.6 Herpetological Surveys

Surveys followed standard protocols [47] and included pitfall traps, standardized transects, and opportunistic searches. Eleven 25cm pitfall traps were installed 10m apart, connected by a 50cm high plastic barrier to guide animals into the traps. Traps were checked twice daily, at 05:30 and 16:30.

Standardized transects were between 200m and 250m and were designed to cover diverse habitats. Point counts were conducted at regular intervals of 10m, during both day (8:00–15:00) and night (18:30 onwards) using headlamps. Transect surveys were supplemented with opportunistic searches which were completed at likely refuges, such as under bark, in leaf axils, and inside hollow logs. Species were identified according to the taxonomy proposed by Glaw & Vences [48].

### 2.7 Inferential Statistical Analysis

Geospatial mapping and statistical analysis were conducted in R (Version 4.4.1)[49]. Open source topographical data at the 1 arc-second resolution was accessed from the NASA SRTM GL1 dataset [50], OpenStreetMap hydrographic layers were accessed from Geofabrik under the Open Database License (https://download.geofabrik.de)[51] and administrative boundaries were obtained from GADM (Global Administrative Areas, v4.1).To standardize survey results, species counts across both categories (eDNA and Visual) were grouped according to their GPS coordinates to allow paired comparisons. Of the 20 eDNA sites, 3 were removed from further analysis, due to failed sequencing (MAK17, MAK20) or sequencing by one primer only (MAK01). As a result, statistical analysis was completed on a total of 17 sites. For these sampling site, data was cleaned, with duplicated species at sites by either sampling type removed, and eDNA classifications of monotypic genera assumed to be the respective species, with all subsequent eDNA analysis completed using distinct OTUs, rather than species classifications to avoid unjustifiable merging of OTUs that mask cryptic species detections, primer-specific biases, and errors in reference databases. We employed this protocol throughout the analysis unless otherwise stated.

To evaluate the species richness at each of the 17 sample sites, we chose the Shannon Diversity Index, instead of other metrics for α-diversity, and used read-count frequencies of distinct OTUs as a proxy for abundance [52], given our data was zero-inflated and contained taxonomically unresolved read counts which may be non-mutually exclusive. OTU read-count was pooled across both the 12S and 16S primers to generate a conservative estimate of species richness, and OTUs detected by both primers were deduplicated to avoid artificial inflation of species richness estimates. Prior to analysis, the dataset was also cleaned to remove ambiguous or non-target sequences, and samples with zero read-counts were excluded. The Shannon Diversity Index (H’) for each sample was computed manually using the formula H’ = -Σ(p_i_ * ln p_i_) where p_i_ is the proportional OTU abundance in each sample [53].

To explore taxonomic primer biases for both the 12S vertebrate and 16S mammal specific primers, we computed the mean detections by both primers across the 17 sample sites for taxa that were detected by both; amphibians, mammals and ray-finned fish. We used distinct OTUs rather than solely sequences resolved to species level to reduce the influence of reference database limitations and to more accurately capture primer-specific amplification bias. Prior to analysis, statistical testing for normality visualised using a Q-Q plot suggested our dataset was non-parametric, therefore we used non-parametric tests for significance in this analysis. Wilcoxon’s Signed Rank tests were used for each taxon to evaluate the significance in differences in detection frequencies between the two primers, which was conducted using the “stats” package in R.

We further analysed the data to visualise the proportion of OTUs resolved to species-level and those that were unresolved to evaluate the effect of GenBank library coverage on effective eDNA detection. Total OTU detections by taxa were computed using both unresolved and resolved sequences and pooled across both the 12S vertebrate and 16S mammal-specific primers, whilst unresolved OTUs were meaned for taxa identified by both primers to provide a conservative estimate of unresolved OTUs, with error bars computed using standard error by the “stats” package in R.

We used a similar approach in defining OTUs by site, to evaluate the efficacy of eDNA in comparison to visual surveying techniques across all taxa. Data cleaning occurred to remove duplicated species at sites detected across both 12S vertebrate and 16S mammal primers, and occasionally by the same primer. The MAK#17 eDNA sample was removed from the analysis due to its exclusion from sequencing, and across taxa the MAK#1 sample was included due to its sequencing by the 16S mammal primer. The total number of OTUs detected by eDNA were calculated as above, as the total number of resolved OTUs and the meaned number of unresolved primers across both the 12S and 16S primers. For both eDNA and visual data, the ‘expected’ number of detections was calculated as the proportion of focal taxa constituting the total inventoried species in the MFPA (n = 316), scaled to the total number of taxa detected by eDNA (n = 83) and visual (n = 125) surveys. The difference to which is displayed in Figure 6. Prior to analysis, statistical testing for normality visualised using a Q-Q plot indicated our dataset was non-parametric, therefore we used the non-parametric Wilcoxon’s Signed Rank test to evaluate levels of significance in difference between our eDNA and visual datasets, conducted by the “stats” package in R.

To estimate total taxon richness in the survey area, and compare the efficacy of eDNA as a survey method compared to conventional techniques, we used Michaelis-Menten functions to fit asymptotes on the observed species accumulation curves (SACs) in the “vegan” package in R, quantifying I_max_ (asymptotic richness) and *K*-values (rate parameter). We selected two classes (amphibians and ray-finned fish), as they offered a sufficient number of taxa detections to allow robust model fitting. First, we selected amphibian detections to evaluate the number of taxa/OTUs in the survey area incorporating both resolved and unresolved OTUs. A Michaelis-Menten model was fitted to the SAC which incorporated 1,000 bootstrapping permutations to generate confidence intervals and estimate asymptotic richness. Secondly, to evaluate the efficacy of eDNA as a survey method in comparison to conventional techniques we used ray-finned fish detections resolved to species-level in both datasets. As before, we fitted a Michaelis-Menton model to the bootstrapped SACs, which incorporated 1,000 bootstrapped permutations to generate confidence intervals, to estimate I_max_and *K*-values. To quantify the survey effort required to achieve a given proportion of asymptotes, we extracted the predicted number of survey hours necessary to detect 95% of the asymptotic richness.

## RESULTS

eDNA analysis of three water samples from each of the 17 sites sampled in the MFPA detected a total of 158 distinct Operational Taxonomic Units (OTUs), 265,154 and 656, 281 reads, using amplification by the vertebrate and mammal specific primers respectively. A comparison of the alpha diversity of each site was inferred using the Shannon Diversity Index calculated from the vertebrate OTU dataset and is visualised in Figure 3 [53]. eDNA metabarcoding analysis detected a mean Shannon Index (*H’*)across sites of 1.58 ± 0.178 with significant variation in species diversity across the 17 sites, with a range of Shannon Index values (*H’*) between 0.379 (MAK#18) to 2.83 (MAK#3), with OTU evenness inferred from the read counts of OTU’s identified to species or genera level.

**Figure 3:**
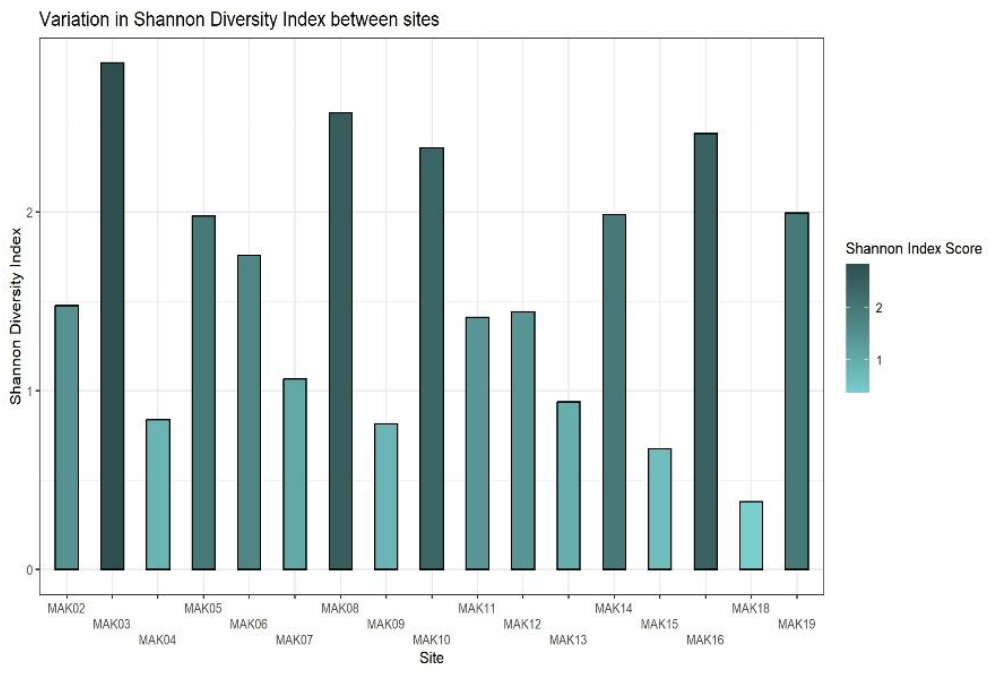
Variation in Shannon Index (H’) between sites detected by eDNA. OTU read-count was used as a proxy for abundance, and distinct (resolved & unresolved) OTUs were pooled across both primers.

Figure 4 indicates the difference in the mean number of distinct OTUs detected using the two primers. A classical Wilcoxon’s Signed Rank Test reported a marginal p-value for ray-finned fish (*W* = 19, *N* = 17, *p* = 0.0890), however paired differences were highly asymmetric, with a permutation-based Wilcoxon-Pratt test reporting a non-significant p-value (*p = 0*.125), suggesting the visual difference is driven by a number of large outliers rather than a consistent effect. Wilcoxon’s Signed Rank tests on the other classes report no significant difference in detections by the vertebrate and mammal primers in both amphibians (*W* = 42, *N* = 17, *p* = 0.187) and mammals (*W* = 28, *N* = 17, *p* = 1.00), although we observed a large difference in median detections of amphibians by the vertebrate primer (3.82 ± 0.456) and the mammal primer (6.44 ± 1.34). The lack of significance in efficacy differences by both primers across taxa suggests that differences in taxa-specific amplification have resulted from a lack of primer specificity.

**Figure 4:**
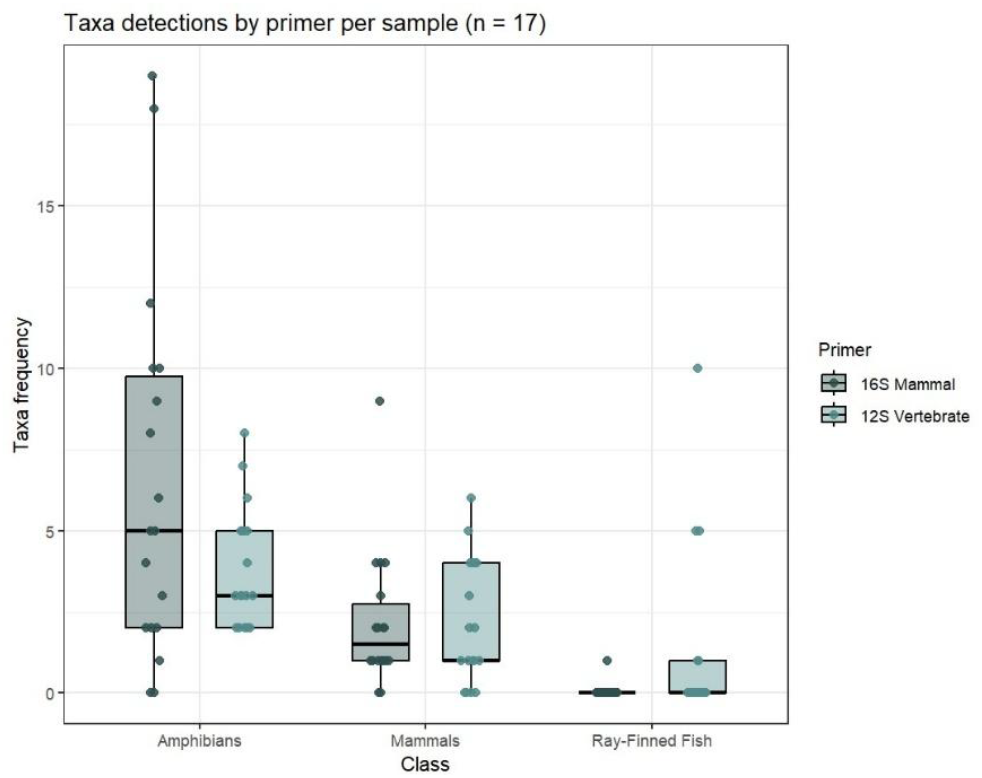
OTUs detected by each primer used, by taxa. Wilcoxon Signed Rank: Amphibians (p > 0.05), Mammals (p > 0.05), Ray-Finned Fish (p > 0.05).

We also found that eDNA analysis of samples detected a significant number of OTUs that were unable to be identified to species level, particularly within amphibians, as displayed in Figure 5. A total of 41 unidentified 12S fragments were detected by the vertebrate primer, and 27 unidentified 16S fragments were detected by the mammal primer. A mean of 20.5 ± 2.5 unidentified amphibian OTUs were observed, alongside 13 unidentified bird OTUs, 3.5 ± 0.5 unidentified mammal OTUs, 3.5 ± 3.5 unidentified ray-finned fish OTUs and 0 unidentified reptile OTUs, averaged across the use of both primers, indicating that a substantial proportion of detections could not be resolved to taxonomic identity in this dataset.

**Figure 5:**
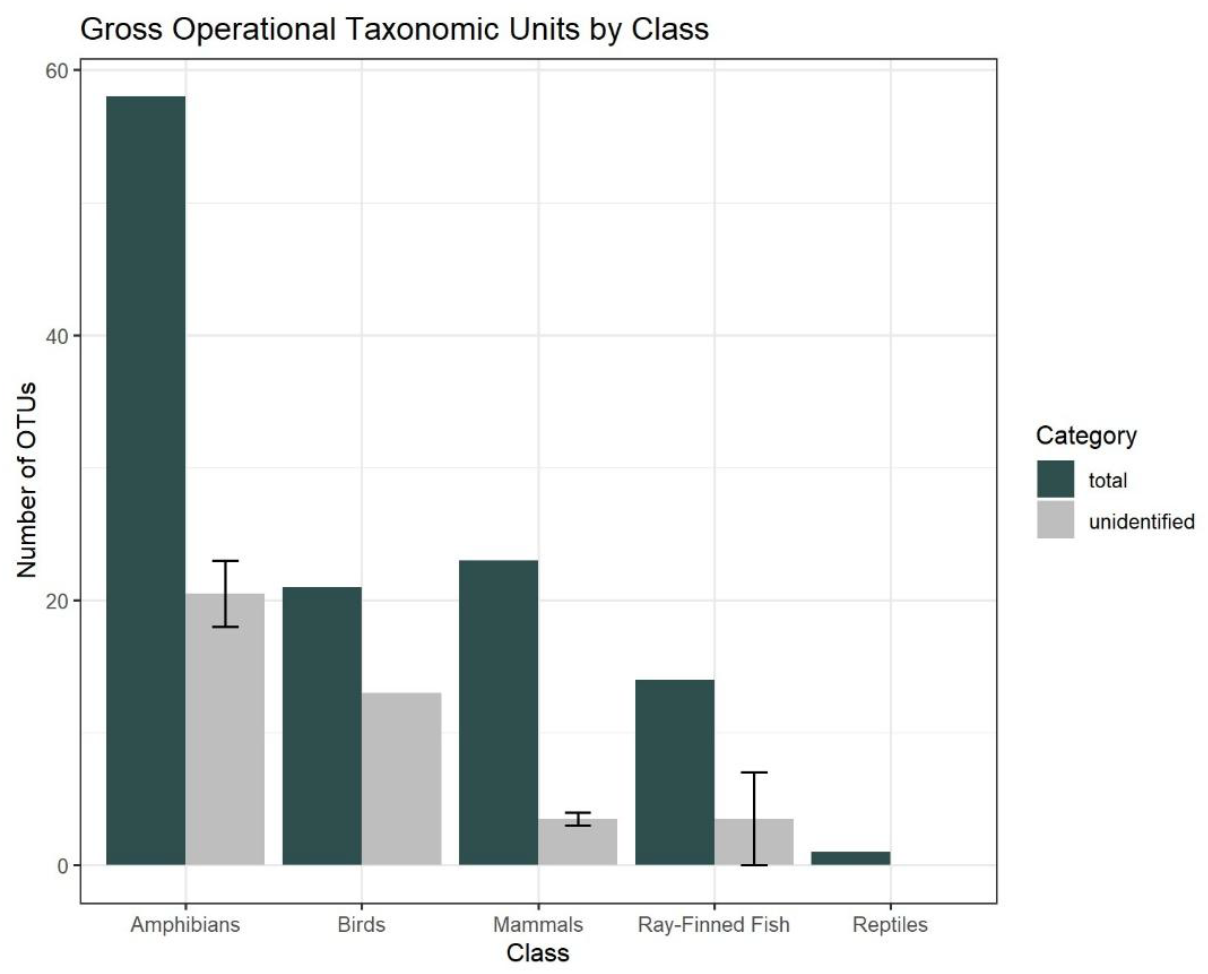
total OTUs (blue), and those unidentified to species level (grey). Error bars are standard error (calculated using both primer datasets for amphibians, mammals, and ray-finned fish).

Figure 6 depicts the mean adjusted difference in species identifications by eDNA sampling and visual techniques from published inventories of known species from the MFPA (n_total_ = 316)[10-13, 54] Expected observations were calculated using the by-class ratios from these inventories and scaled to the observed number of taxa detected by our surveys (visual n_total_= 125, eDNA n_total_ = 83), with the difference between observed and expected observations presented, by taxa, in Figure 6 with the zero line indicating 100% efficacy, to give an indication of sampling bias by taxa. Sample sites are treated as pseudo-repeats. Analysis using the Wilcoxon Signed Rank test indicate that eDNA metabarcoding is therefore significantly more effective at detecting all taxa analysed in this study. eDNA was more effective at detecting amphibian taxa (*W* = 0, *N* = 15, *p* = 0.000725); with eDNA detecting a median difference of -11.5 ± 1.03 species, whilst visual surveys detected a median difference of -24.9 ± 0.696 species in comparison to the scaled inventoried taxa ratios for the MFPA. eDNA sampling was also more effective at detecting reptile taxa (*W* = 0, *N* = 14, *p* = 0.000877), with eDNA a median difference of -15.6 ± 0.0762, in comparison to visual surveys (-21.9 ± 0.294), from published inventories. Ray-finned fish detections were also significantly higher for eDNA than conventional surveys (*W* = 1.00, *N* = 10, *p* = 0.00788), with eDNA detecting a median difference of -3.84 ± 0.405 species, in comparison to visual surveys (-6.42 ± 0.3). eDNA was also significantly more effective at detecting birds than visual surveys were (*W* = 6, *N* = 18, *p* = 0.000107). eDNA surveys detected a median difference of - 27.9 ± 0.448 species, compared to -34.5 ± 1.34 species by visual surveys, from the scaled inventoried taxa of Makira. eDNA was also significantly more effective at detecting mammals than visual surveys (*W =* 0, *N* = 15, *p* = 0.000718). eDNA detected a lower median difference of -12.8 ± 0.48 in comparison to visual surveys (-21.9 ± 0.00). When considered together, the use of eDNA increased overall species detection rates by 66.4% in comparison to the use of visual surveys alone, underscoring the complementarity of the two methods.

**Figure 6:**
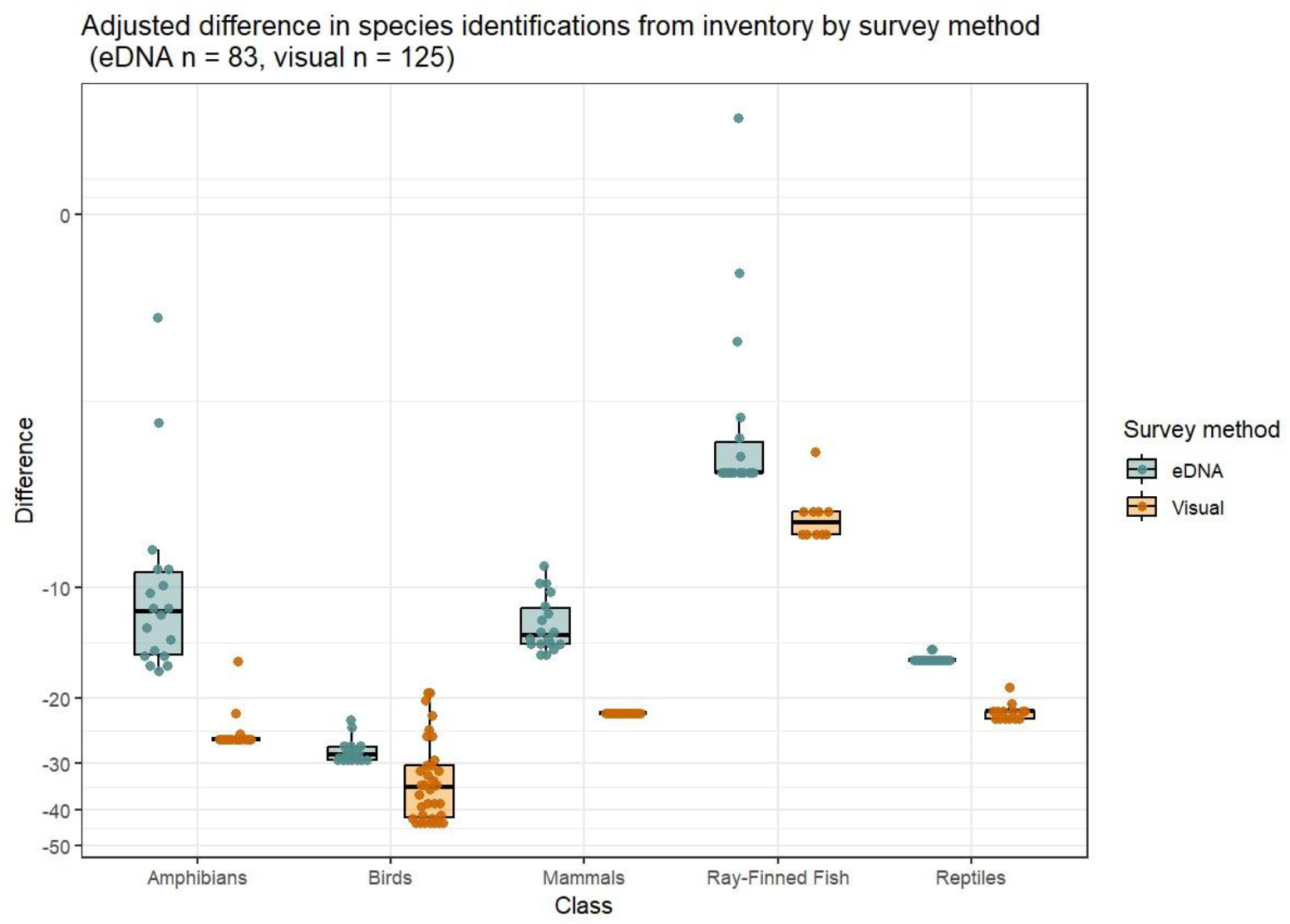
Adjusted difference between total observed and expected species identifications (n = 316), standardized to sample size, by survey method. Wilcoxon Signed Rank: Mammals (p <0.01), Amphibians (p < 0.001), Reptiles (p < 0.001) Ray-Finned Fish (p < 0.01) Birds (p < 0.001); Y axis is pseudo log transformed.

Figure 7 evaluates the efficacy of eDNA versus conventional ichthyological surveys in detecting ray-finned fish in the MFPA. eDNA detections were conservatively defined as OTUs identified to species level, and Figure 7 presents the bootstrapped species accumulation curves of both eDNA and conventional methods, to which we fitted a Michaelis-Menten model, to estimate I_max_ and *K*-values. When standardised to cumulative person hours, eDNA sampling detected species at a rate of 0.41/hour compared to 0.03/hour using conventional survey techniques, indicating an almost 15-fold increase in efficiency from our sampling. The Michaelis-Menten model fitted to the SAC estimated the I_max_ of eDNA detections, which reached 23.3 species, a value close (129%) to the known inventoried ray-finned fish of the MFPA (18) [13]. On the other hand, the modelled asymptote of the conventional survey reached 9.2, just 51% of the inventoried species list. We additionally report that the Michaelis-Menten model for eDNA estimated a *K*-value of 7.05, and that 134 cumulative person hours would be required to detect 95% of asymptotic richness, whilst the extrapolated Michaelis-Menten model for conventional surveying estimated a *K*-value of 69.8; requiring 1,327 cumulative person hours to detect 95% of the asymptotic richness, itself just 51% of the known richness of the MFPA. This indicates eDNA’s had a 10.0 fold increase in efficiency in detecting ray-finned fish in comparison to conventional methods in this survey.

**Figure 7:**
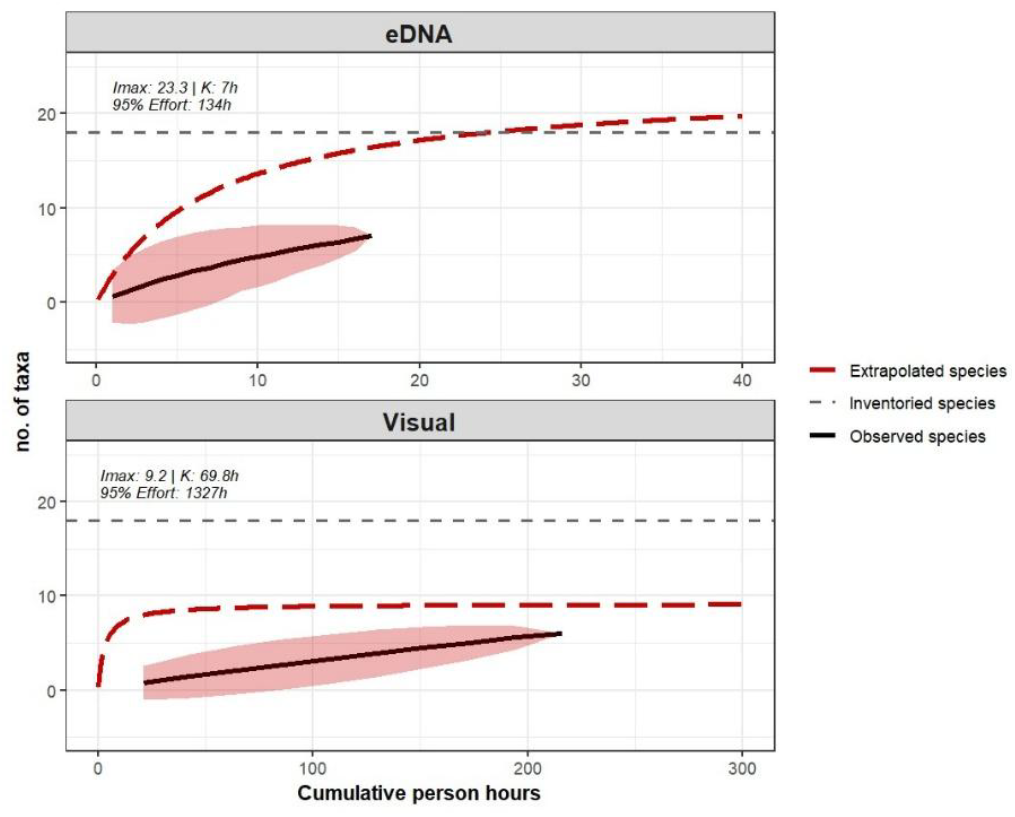
Bootstrapped species accumulation curves for eDNA (top) and conventional surveys (bottom) of ray-finned fish. The detections of species were bootstrapped 1000 times to generate confidence intervals (red shading). A Michaelis-Menten model was fitted to the SAC and used to estimate asymptotic richness (red dashed-line). Total inventoried ray-finned fish species of the MFPA (18) displayed for comparison (grey).

The result presented in Figure 5 is further supported by Figure 8 which presents bootstrapped species accumulation curves for the eDNA detection of amphibians, across our 17 samples, comparing the efficacy of defining taxa using a resolved-to-species approach, or an unresolved OTU approach during analysis. The Michaelis-Menten model estimated an asymptote of 33.5 when using a resolved-to-species approach, a value close to the 21 sequenced species present in the MFPA, but 2.0 times lower than the MFPA amphibian species checklist [11]. On the other hand, if using all distinct OTUs detected by both primers, the modelled asymptote was much higher than the regional checklist (68) at

**Figure 8:**
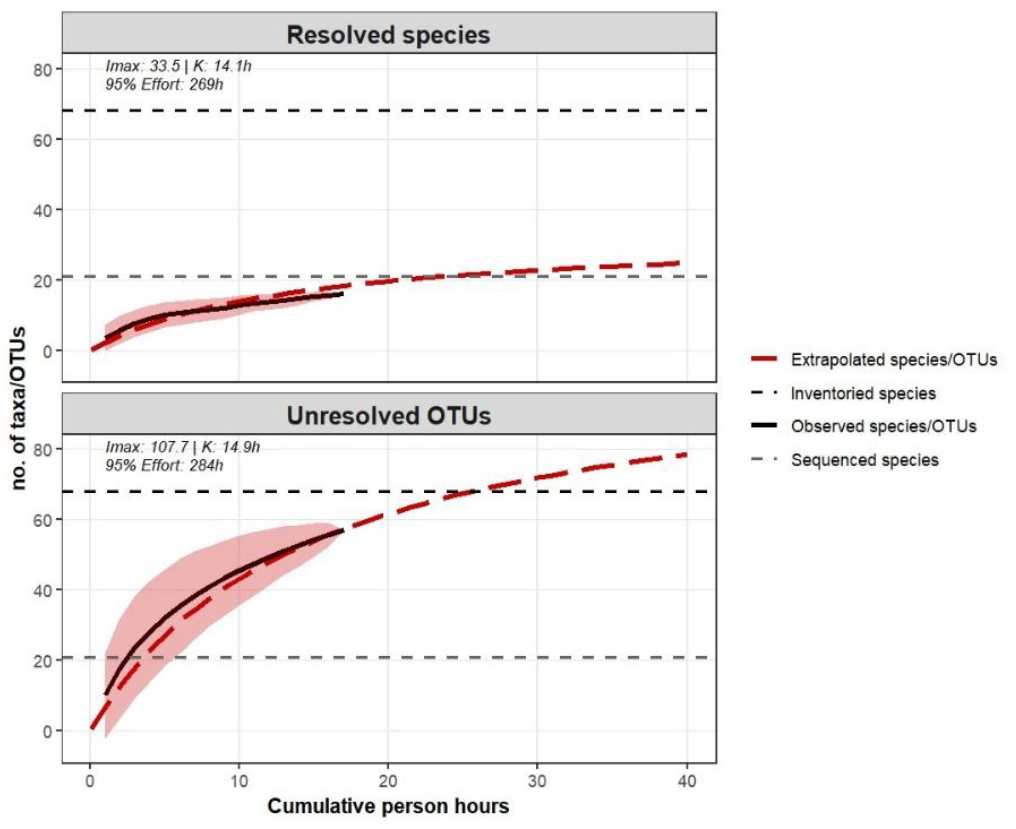
Bootstrapped species accumulation curves for eDNA amphibian detections using resolved-to-species (top) or unresolved OTU approaches (bottom). The detections of species and unique OTUs (black) were bootstrapped 1000 times to generate confidence intervals (red shading). The asymptotes (red dashed-line) were modelled using a fitted Michaelis-Menten model. Total inventoried amphibian species (68) from MFPA (black), and amphibian species sequenced and available in GenBank (21) from MFPA (grey) for comparison.

107.7 species, indicating that a substantial proportion of amphibian diversity in the MFPA remains taxonomically unresolved in current GenBank databases, and conservatively defining eDNA detections as resolved-to-species OTUs, will significantly underestimate taxonomic diversity. However, we report that both extrapolated SAC models had similar *K*-values of 14.1 and 14.9 hours, with estimated effort to reach 95% of asymptotic richness being 269 hours and 284 hours for assigned species and un-assigned OTUs respectively, indicating significant survey effort is required for significant species richness detection.

## DISCUSSION

### Comparative Effectiveness of eDNA vs. Conventional Surveys

Efficacy evaluation of our eDNA metabarcoding protocol in direct comparison to conventional survey methods, quantified using gross species detections, observed significant taxa-specific efficacy variation, with eDNA displaying a higher efficacy than conventional survey techniques across all taxa (Figure 6). The analysis revealed that eDNA metabarcoding was significantly more effective than traditional transect surveys at capturing bird, amphibian, reptile, fish and mammal biodiversity. The higher efficacy result for ray-finned fish was further supported by extrapolations of species accumulation curves (Figure 7), which highlighted a much more accurate reflection of inventoried species in the MFPA by the extrapolated asymptotes of eDNA analysis in comparison to conventional electrofishing and seine netting. Our finding that eDNA metabarcoding is more effective at detecting herpetofauna and ray-finned fish specifically is consistent with previous research [55-57], although our result for other taxa such as birds and mammals may be due to weaknesses in conventional survey design on this project. These findings highlight the particular strength of eDNA in sampling cryptic or elusive taxa, especially in environments where conventional techniques are likely to under-detect true species presence.

Additionally, we observed sampling bias in conventional sampling techniques due to variation in sampling effort. However, this outperformance in capturing diversity by eDNA is due to a number of issues with conventional survey techniques, primarily a high frequency of false negatives. False negatives in conventional survey techniques are elevated by a number of environment-level and species-level characteristics. Environment-level challenges include habitat accessibility, weather variability, and climatic variables [58], whilst species-level challenges associated with conventional survey techniques include behavioural attributes, difficulty in identifying genetically and visual cryptic species, and low population densities, all of which eDNA metabarcoding analysis is more robust to detect [22, 59, 60]. Integrating eDNA with conventional survey techniques increased species detection rates by 66.4%, demonstrating that these methods are not necessarily interchangeable but highly complementary. Together, they provide a more comprehensive picture of biodiversity, particularly in complex, under-sampled ecosystems like the MFPA.

Species accumulation curves for ray-finned fishes showed that both our eDNA and visual sampling did not reach an asymptote suggesting that additional effort would yield new species detections. This finding reinforces the value of continued sampling for tropical freshwater fishes in complex environments like the Makira plateau. Michaelis-Menten models fit to these SACs revealed that the modelled asymptote for conventional survey techniques did not reach the number of known freshwater fish species in the MFPA, suggesting that these surveys were unsuccessful in effectively capturing species richness, likely due to insufficient sampling effort and low efficacy. We were unable to generate comparable curves for other taxa due to insufficient site-specific data on conventional survey effort, which was not tracked at the level of robust individual sites. As this was a multi-taxa rapid inventory, we nonetheless used all available detection data from these groups to assess relative survey effectiveness and biodiversity patterns.

However, our species accumulation curves of amphibians presented in Figure 8, indicate that treating unresolved OTUs as distinct taxons, yielded a more accurate approximation of the total described species in the national park. Conversely, evaluating resolved OTUs significantly underestimated the total alpha richness of the site. This conclusion suggests that whilst many papers conservatively exclude unresolved OTUs for ecological analyses, such approaches may acutely underestimate and potentially compromise the accuracy of biodiversity assessments, that feed conservation planning.

Therefore, whilst eDNA may in some cases outperform conventional survey methods, the treatment of unresolved or resolved OTUs may skew richness estimates, with current conservative practices being inherently flawed.

### Site-Specific Variation in Species Diversity

Detected differences in the Shannon Diversity Index, as a measure of alpha diversity, of almost ten-fold, quantified using the number of distinct OTUs and their read-counts from each sample site supported our expectation of significant site-specific differences in diversity. However these results must be interpreted with caution given sample sites were located within the same watershed system, and differences in alpha diversity are unlikely to be due to high true beta diversity. Using read-counts as a proxy for abundance is a known drawback of eDNA data, as the relationship between DNA concentration and organism abundance is poorly understood. Whilst recent research has successfully estimated abundance of target species in marine ecosystems from eDNA read counts [52, 61], many of these studies remain limited to mesocosm experiments and read-count number from surveys in natural environments poorly explain abundance[62, 63] although some success has been observed [64].

In lotic environments, eDNA concentrations exhibit high spatiotemporal variability as eDNA moves downstream from the point source in a complex plume [65, 66], due to a number of highly stochastic environmental and biological variables known to effect DNA shedding and degradation. Spatial eDNA heterogeneity in freshwater ecosystems is linked to environmental factors affecting eDNA dispersion including water flow conditions, temporal variation, and vegetation structure [67, 68] which varied considerably across the 17 sites we sampled (see Figure 1C-E). Additionally, eDNA degradation is a major factor affecting detectability and curtails the power of spatial and temporal inferences drawn from analysis [69]. Degradation is affected by a number of variables including UV-B levels, water pH, and temperature [70], measurements of which should be incorporated into reporting of eDNA sampling to allow environmentally driven variation to be evaluated. Furthermore, the presence of naturally occurring proteases and compounds that reduce the accuracy of eDNA analysis by interfering with PCR analysis, that includes tannins, humic acids, heme, and other enzymatic inhibitors significantly impact eDNA detection efficacy [71, 72]. To mitigate this, we used an inhibitor removal kit to purify samples prior to PCR amplification and minimise inhibition in our samples, alongside a blocking primer to minimise non-target DNA amplification, and would encourage incorporation of inhibitor and non-target nucleotide removal techniques during eDNA refinement processes. Our finding of significant variation in alpha diversity across sites with similar macro-ecological characteristics suggest that understanding variables leading to spatial heterogeneity in eDNA dispersion and degradation are essential in refining models for, upstream species localisation and abundance inference, particularly in the context of cryptic and rare species.

### Taxonomic Resolution and Data Gaps in Reference Libraries

A key limitation observed in this study was the high number of unidentified OTUs, particularly in amphibians. While eDNA metabarcoding offers a promising approach for species detection, its efficacy is ultimately constrained by the completeness and quality of reference databases [73]. In this study, a substantial proportion of OTUs could not be assigned beyond higher taxonomic levels, due to poor representation of Malagasy taxa in GenBank. Such gaps in reference libraries hinder the resolving power of PCR analysis, and additionally the identification of potentially new species. For example, we detected two distinct sequences belonging to the monotypic *Galidia* mongoose genus, with the unresolved sequence containing two point mutations in the target region, suggesting issues with reference libraries or a possible undescribed species detection. In comparison, the 16S mammal-specific primer duplicated detections of both *Boophis englaenderi* and *Guibemantis liber* with small point-mutation differences between the sequences and resolved them to species level. Evaluating possible undescribed species detections and duplicated resolved species detections is made significantly more challenging by poor reference libraries.

Addressing these challenges will require targeted efforts to expand reference libraries, including voucher-based sequencing of Malagasy vertebrates. Historically, extracting DNA from museum specimens has been challenging due to degradation over time. However, recent advancements in DNA extraction and sequencing technologies have significantly improved the recovery of genetic material from such samples [74, 75]. These developments enhance the potential to enrich reference databases with sequences from archived specimens, thereby reducing the number of unidentified OTUs in understudied regions and taxa. Taxonomic verification using integrative approaches - such as combining eDNA with conventional morphological identification and targeted sequencing will also be critical to improving the accuracy and reliability of future biodiversity assessments [76].

Further comparative analyses of OTU assignment across primers could have provided additional insights into amplification bias and taxonomic resolution [77]. However, logistical and capacity constraints prevented these analyses from being conducted during the study period, limiting our ability to assess primer-specific performance comprehensively. As noted in prior studies, primer selection significantly affects detection accuracy, particularly in taxa with poor reference representation [78]. Pre-collection agreement of appropriate primers, aligned with the goals and outputs of a study, is critical to maximizing detection success and ensuring comprehensive taxonomic coverage and interpretable results.

### Primer Specificity and Amplification Bias

We found significant differences in mean species detection by taxonomic group between the 12S vertebrate specific and 16S mammal specific primers used for amplification of OTUs across all the samples analysed (Figure 4) (see [79] for primer design). Both primers displayed a significantly higher mean species per site detection in Amphibia, followed by Mammalia and finally Actinopterygii, likely due to a combination of differences in intrinsic taxa diversity between classes, and taxa-specific differences in behavioural interactions with lotic habitats. Evaluating the efficacy of eDNA metabarcoding of freshwater sampling in lotic environments for non-aquatic taxa should therefore be a future research priority as detection of non-aquatic target taxa may suffer from disproportionate reduced detection efficacy with ramifications for survey outputs, specifically abundance estimates. Our findings also suggest that differences in taxa-specific amplification are as a direct result of issues with primer specificity and design. We found no significant difference observed in mean species detection across Amphibia, Mammalia and Actinopterygii between the vertebrate and mammal-specific primers. Specifically, a higher mean detection of amphibians across sites by the mammal-specific primer rather than the vertebrate primer raises significant concerns of amplification of non-target OTUs and PCR bias. Primer refinement and redesign clearly needs to occur to mitigate PCR bias which is commonly a result of PCR drift, interspecific variation in gene copy, denaturation efficiency and primer binding affinity [80-82]. Amplification using two primers here suggests primer binding affinity as the most significant factor driving taxa-specific amplification differences, therefore we suggest the inclusion of positive controls with known OTU fragments to examine primer-specific variation in amplification. Moreover, downstream PCR replicate sequencing reported two duplicate OTUs assigned to species level. *Boophis englaenderi* was recorded as separate OTUs by the mammal-specific primer, whilst *Guibemantis liber* was recorded as separate OTUs with a difference of one single nucleotide polymorphism by the vertebrate primer. These records indicate flaws in the post-sequencing workflow which included demultiplexing, denoising and dereplication. Combined with issues of primer specificity and PCR bias, which together pose significant challenges in reliably surveying taxonomic groups, the importance of combining OTUs detected by more than one primer into a meaningful single value for species detections cannot be overstated, and whilst having precedent often over or under-estimates species richness, the robustness of which increases with improved coverage in reference databases [77].

### Challenges and Considerations in eDNA Sampling

A key consideration in the use of eDNA metabarcoding remains inferring conservation-relevant biological metrics from eDNA data, such as effective population size, census size, and biomass due to the lack of explanatory power between read-count number and abundance. This is due to the high stochasticity of environmental and biological factors affecting DNA shedding and degradation. Unidentified OTUs may reflect a combination of incomplete reference databases, DNA degradation leading to reduced sequence quality or length, and marker-specific limitations, suggesting that additional loci or complementary primers may improve taxonomic resolution in future surveys.

Because this was a rapid, multi-taxa biodiversity inventory covering fish, birds, mammals, amphibians, and reptiles, no single eDNA sampling method could be entirely effective across all taxa.Aquatic eDNA sampling, while well-suited for detecting freshwater species, has inherent limitations when applied to terrestrial and fossorial organisms. Our results demonstrated the effectiveness of water-based sampling for certain groups, as eDNA significantly outperformed conventional surveys in detecting amphibians, and ray-finned fish. However, whilst the results in Figure 6 present scaled detections, gross detections by eDNA per site for the terrestrial vertebrate groups (mammals, birds and reptiles), were more infrequent than the aquatic groups. This aligns with existing research indicating that terrestrial reptiles shed minimal DNA and have limited interaction with water sources, making aquatic eDNA an unreliable detection method for this group [70].

To address these limitations, future multi-taxa biodiversity surveys in diverse tropical ecosystems should incorporate complementary eDNA sampling strategies tailored to taxon-specific needs. Novel eDNA technologies are rapidly evolving, offering new opportunities to overcome the limitations observed in the MFPA and improve biodiversity detection across multiple taxa. Automated and multi-sampler eDNA technologies could further enhance future surveys by increasing sampling efficiency, reducing human error, and allowing for long-term, continuous sample collection. Autonomous samplers, such as the DOT-NM Autosampler, can collect multiple eDNA samples at pre-programmed intervals, ensuring a more representative snapshot of biodiversity across different times of day and environmental conditions (NatureMetrics; 1 Occam Court, Surrey Research Park, Guildford, GU2 7HJ United Kingdom, https://www.naturemetrics.com/dot-nm-autosampler). For terrestrial reptiles, soil eDNA sampling and substrate swabbing at basking sites, burrows, or movement corridors would likely yield better results, as reptiles shed DNA primarily through skin and faecal deposits rather than into water [83]. This approach would be particularly beneficial for fossorial species, whose limited surface activity makes them difficult to detect through conventional methods. Rainwater eDNA collection could offer an effective, minimally invasive method for detecting arboreal and ground-dwelling species by capturing DNA washed from the canopy and forest floor during precipitation events. Airborne eDNA sampling, using HEPA filters or drone-mounted vacuum collection, has the potential to significantly enhance the detection of birds and mammals, particularly those that roost or nest in dense forest canopies, where direct observation and physical sampling are challenging [84].

Additionally, leaf swabbing could be a valuable tool for detecting species that interact with vegetation, especially in areas where terrestrial mammals and reptiles frequently climb or feed. As eDNA methodologies continue to advance, these emerging approaches represent some of the most promising solutions for improving biodiversity assessments, expanding detection capabilities across a wider range of taxa while minimizing disturbance to the ecosystem.

### Future Research Directions

While eDNA is an efficient and cost-effective tool for detecting rare and cryptic species, it rarely provides insights into behaviour, population dynamics, or habitat use. For more comprehensive biodiversity assessments, future efforts should consider integrating eDNA with conventional surveys, as the two methods offer complementary strengths and can compensate for each other’s limitations [85]. Further, combining eDNA with other non-invasive technologies, such as camera trapping, bioacoustics, 3D scanning, and drone surveys-can enhance conservation monitoring, particularly in dense forest environments where species detection is inherently challenging.

Community-based conservation initiatives are critical for ensuring biodiversity data remains accessible and beneficial to local stakeholders. Incorporating eDNA sampling into local monitoring programs fosters consistent, participatory biodiversity assessments, strengthening conservation efforts through direct community engagement. Training local conservation teams to collect, process, and analyse eDNA data enhances long-term ecological monitoring and empowers communities to take an active role in protecting their environment [86]. In the MFPA, integrating eDNA into local conservation efforts could improve species detection, track environmental changes more effectively, and provide communities with the tools to respond to emerging threats.

Tropical rainforest ecosystems like Makira-despite their size and ecological significance-are increasingly threatened by a convergence of pressures common across much of the global tropics. During this expedition, we documented widespread anthropogenic impacts, including deforestation near Sahametreha, gold and gemstone mining, rice paddy expansion, and watercourse diversion. Free-ranging livestock (pigs and cattle) were found deep within forested areas, and eDNA analysis confirmed the presence of multiple invasive species (e.g., *Channa maculata, Rattus rattus, Gambusia holbrooki*). Signs of poaching, including bird and lemur traps and reports of armed individuals, further underscored the vulnerability of even the most remote and protected, forested regions. These challenges are by no means unique to Madagascan forests, and are seen in tropical forests across Southeast Asia, the Amazon, and Central Africa. These compound threats highlight the urgent need for scalable, non-invasive biodiversity monitoring tools. Integrating eDNA with conventional survey methods offers one such approach, enabling broader, more effective, tracking of biodiversity at the community-level across threatened tropical systems.

## Notes

https://doi.org/10.7910/DVN/QK3E1D

### Competing Interest Statement

Christina Biggs at the time of project execution was an employee of Re:wild, Cosmo Le Breton is a founder of the RIDGES Foundation. Both organisations provided funding to this expedition. The funders had no role in study design, data collection, analysis, interpretation, or the decision to publish.

### Summary of Updates

Clarity of methods section significantly enhanced, cosmetics of figures improved.

